# Cell elimination strategies upon identity switch via modulation of *apterous* in *Drosophila* wing disc

**DOI:** 10.1101/674168

**Authors:** Olga Klipa, Fisun Hamaratoglu

## Abstract

The ability to establish spatial organization is an essential feature of any developing tissue and is achieved through well-defined rules of cell-cell communication. Maintenance of this organization requires elimination of cells with inappropriate positional identity, a poorly understood phenomenon. Here we studied mechanisms regulating cell elimination in the context of a growing tissue, the *Drosophila* wing disc and its dorsal determinant Apterous. Systematic analysis of *apterous* mutant clones along with their twin spots shows that they are eliminated from the dorsal compartment via three different mechanisms: sorting to the ventral compartment, basal extrusion, and death, depending on the position of the clone in the wing disc. We find that basal extrusion is the main elimination mechanism in the hinge, whereas apoptosis dominates in the pouch and in the notum. In the absence of apoptosis, extrusion takes over to ensure clearance in all regions. Notably, clones in the hinge grow larger than those in the pouch, emphasizing spatial differences. Mechanistically, we find that limiting cell division within the clones does not prevent their extrusion. Indeed, even clones of one or two cells can be extruded basally, demonstrating that the clone size is not the main determinant of the elimination mechanism to be used. Overall, we revealed three elimination mechanisms and their spatial biases for preserving pattern in a growing organ.

## INTRODUCTION

Multicellularity requires precise spatial organization of cells during development. Aberrant cells that arise as a result of sporadic mutations or chromatin defects can challenge the robustness of a developmental program. For example, the cells that acquire incorrect positional identity disrupt the proper spatial organization. Those cells are potentially dangerous and cannot be tolerated. Therefore, mechanisms that ensure elimination of such cells are in place, yet they remain poorly understood. Arguably, one of the best-characterized systems to study the spatial organization of a developing tissue is the *Drosophila* wing imaginal disc. This organ is amenable to mosaic technique, which is particularly useful for studying interactions of differently specified cells *in vivo*. The pattern in the wing disc is set by restricted expressions of fate determinants that are turned on in a sequential manner. Engrailed and Apterous (Ap) define posterior and dorsal fates, respectively, and lead to the formation of compartment boundaries that provide lineage restriction (1–3). Further subdivisions are achieved by restricted expressions of Vestigial in the pouch, Homothorax and Teashirt in the hinge and the Iroquois complex in the notum (4–8). The cell clones with altered expression of such genes disrupt the tissue pattern and trigger a set of common events. Such clones round up to minimize contact with their neighbors. Some of the clones were reported to undergo apoptosis or bulge out of the tissue forming cysts (9–14).

In addition to separating opposing compartments from each other and preventing cell mixing, the compartment boundaries also act as signaling centers. The morphogens Decapentaplegic (Dpp) and Wingless (Wg), secreted from these centers, form concentration gradients and orchestrate proper tissue size and shape (15–19). Disruption of morphogen gradients also triggers a set of common events that will eventually restore the pattern via elimination of the mis-positioned cells (9, 10, 13, 20–23). Strikingly, in all these cases there is a strong regional bias. For example, *thickveins* mutant cell clones, that lack the Dpp receptor, undergo apoptosis or basal extrusion in the medial wing disc where the pathway activity is high, yet they can be tolerated laterally where the pathway activity is naturally low (21, 24, 25). Therefore, disruption of the pattern in the tissue prompts the mechanisms in place to restore it.

Here, we aimed to understand how cells with altered expression of the dorsal determinant Ap are eliminated from the tissue. It has been shown that Ap expressing clones are eliminated from the ventral compartment; whereas clones expressing Ap inhibitor *Drosophila* LIM only (dLMO) are eliminated from the dorsal one (11). Ap acts as a transcription factor (26). Via regulation of its target genes glycosyltransferase Fringe and Notch ligand Serrate in dorsal compartment it mediates activation of Notch and Wg signaling pathways at the D/V compartment boundary (27–31). Similar signaling was observed around Ap or dLMO cell clones if they happen to be in the incorrect place (11, 32). Depletion of Notch signaling within the clones, using *Notch^DN^*, was shown to prevent almost all elimination in the pouch region (11). The same rescue effect was observed when the apoptosis of clones was prevented by expression of *p35* (11). This highlights the importance of the ectopic signaling for the elimination process and defines apoptosis as its main executor.

Notably, all these experiments were focused on the clones located in the pouch. However, whether clones in other regions behave the same remains unknown. Here we take a quantitative approach to define the contributions of different strategies – apoptosis, basal extrusion and sorting – employed by the tissue to deal with mis-positioned cell clones. Importantly, we did not limit our analysis to a specific region, and characterized *ap* clone behavior throughout the tissue. Our approach revealed a striking regional bias of the contributing mechanisms.

## RESULTS

### A new *apterous* allele and use of twin-spots allow systematic and quantitative analysis of clone elimination

Systematic analysis of the behavior of *ap* mutant clones has not been possible until recently due to technical limitations of classical clonal analysis approaches. This is because the *ap* locus lies between the centromere and the canonical flippase recognition target (FRT) site on the right arm of the second chromosome. Hence this FRT site cannot be used to generate *ap* mutant patches. To circumvent this problem, we used a new *ap* allele generated by Bieli et al. (33, 34), where a well-positioned FRT (f00878) was used to generate a deletion of the whole coding region of *ap (ap^DG8^)*. Using this new tool, we first generated positively marked *ap* mutant clones as well as clones with ectopic Ap expression. The wild-type control clones were distributed uniformly throughout the disc (Fig 1A). In contrast, clones altered for Ap function displayed compartment bias. In agreement with former reports (11, 32), cells that lose *ap* expression are underrepresented in the dorsal compartment (Fig 1B) and, likewise, cells with ectopic Ap expression are eliminated from the ventral compartment (Fig 1C). The misspecified clones that remained in the tissue minimize their contact with surrounding native cells and display ectopic boundary signaling (detected by Wg) at the clone border, where Ap-expressing and Ap-non-expressing cells contact each other (Fig 1B, arrows). Thus, cells are cleared from the region where they do not normally belong, pointing at the existence of intrinsic mechanisms that detect and get rid of misspecified cells and hence contribute to the maintenance of compartment organization.

**Fig 1.**
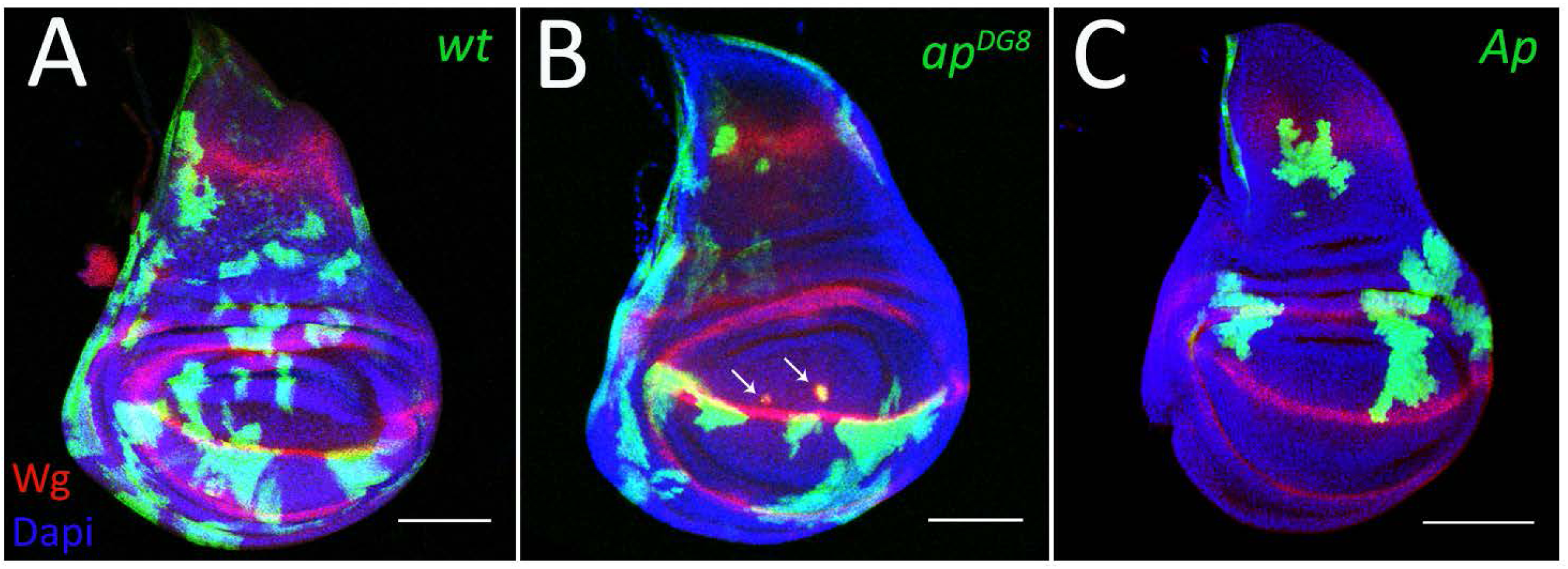
The recovery of clones altered for Ap function is compartment biased. (A-B) Third instar wing discs with GFP-marked wild-type (A), *ap^DG8^* (B) and Ap-expressing (C) clones. The arrows point to ectopic Wg expression around mis-specified clones. Scale bars represent 100µm. Hereinafter disc orientation is dorsal is up, anterior is to the left.

In order to have a disc intrinsic measure of how many clones were originally generated we utilized a classical mitotic recombination approach that allows to generate mutant clones together with their wild-type twin sisters. To mark mutant clones positively we placed *GFP* on the chromosome that carried *ap* mutation. Therefore, in our set-up the *ap* homozygous mutant clones were marked positively by two copies of *GFP*, whereas their sister wild-type clones (twin spots) - by the absence of *GFP*. To understand what happens to the mutant cells after their induction, we followed their fate in a time-course experiment using this set-up. We induced clones shortly before D/V boundary formation, at 46h after egg laying (AEL), and took time points every 10h (Fig 2A). The dorsal clones of each time-point were categorized into 3 groups (Fig 2B). The first group includes pairs of *ap* mutant and wild-type clones (Fig 2B (a)). This group reflects the number of mutant clones that remained in the dorsal compartment at a particular time point. The second group contains wild-type clones without their mutant sisters (Fig 2B (b)) and corresponds to the number of *ap* mutant clones which had already been eliminated. The last group includes wild-type clones in the dorsal compartment that have their mutant twins in the ventral part (Fig 2B (c)), suggesting that those mutant clones are presumably clones of a dorsal origin that had been sorted to the ventral compartment. The percentages of clone pairs in each group normalized to the number of all dorsally located wild-type clones are shown (Fig 2G-I, red lines).

**Fig 2.**
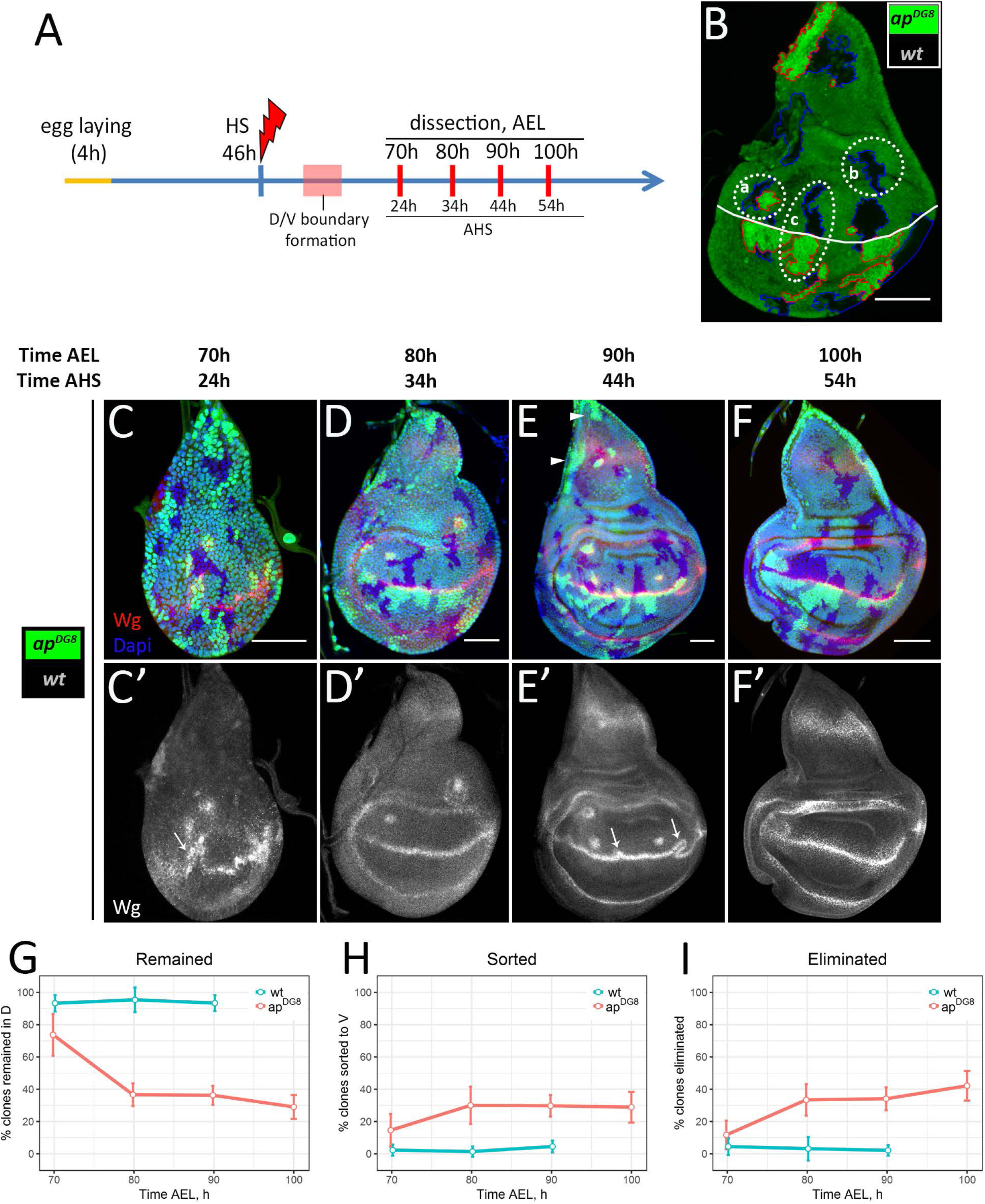
The dynamics of clone elimination. (A) Time-course scheme. (B) Strategy of clonal analysis. Example of the disc with *ap* mutant (2 copies of GFP, red outline) and wild-type sister (absence of GFP, blue outline) clones. (a) – wild-type clone together with the mutant twin; (b) – wild-type clone without the mutant sister; (c) – wild-type clone with *ap* mutant sister in the opposite compartment. White line corresponds to the D/V boundary. (C-F) Wing discs of indicated times containing differently marked wild-type and *ap^DG8^* sister clones. (C’-F’) Wg channel of C-F. (G-I) Plots represent amount of dorsal clones that remained in the dorsal part (G), have been sorted to the ventral part (H) or completely eliminated from the disc (I) as a function of time. Number of clones in each group was normalized to the number of dorsally located wild-type clones (per disc). Red lines correspond to the ratios of *ap^DG8^* clones to their wt sisters (experimental discs); blue lines correspond to the ratios of wt clones to their wt sisters (control discs, shown on Fig S1). Note, the control discs were analysed only at 70h, 80h and 90h AEL. At least 15 discs were analysed for each time-point. Data represent mean±CI (95%). Scale bars represent 50µm.

At 24h AHS (the first time-point), almost 75% of dorsally located wild-type clones had their mutant twins (Fig 2C, 2G). Interestingly, the clones and their sisters had similar sizes (about 8-12 cells) at that time point. This indicates that the mutant cells initially were able to grow in the dorsal compartment. However, 10 hours later (34h AHS) the amount of *ap* clones recovered in the dorsal part dropped sharply (Fig 2D, 2G). Less than 40% of mutant clones remained in the dorsal disc. Those clones were much smaller and more circular compared to their wild-type sisters. In the next 20 hours, the number of dorsally located mutant clones declined only slightly (Fig 2E-F, 2G). Nearly 30% of mutant clones had remained in the dorsal disc at 54h AHS (the last time point). Interestingly, the clones at the very proximal notum (disc tip) and lateral notum regions were not eliminated (Fig 2E, arrowheads).

The mutant clones that were removed from the dorsal compartment had been either sorted to the ventral one (Fig 2H) or eliminated from the disc tissue completely (Fig 2I). We observed relatively high number (15%) of dorsally located wild-type clones with their mutant sisters in the ventral compartment at the first time-point (70h AEL) (Fig 2C, 2H). The number of such clones nearly doubled at the second time-point (80h AEL) (Fig 2D, 2H). The sorted mutant cell clones accumulated at the D/V boundary from the ventral side (visible in Fig 2D-F). Clone sorting is coupled to boundary reorganization (Fig 2C’, 2E’, arrows). We observed that the ectopic signaling induced between the mutant cells and the surrounding Ap-expressing cells can be incorporated into the regular compartment boundary if a mutant clone happens to arise in close proximity to the D/V boundary. Importantly, the deformed D/V boundary straightens after the sorting has been completed, as we nearly never observed boundary deformations at 100h and later (Fig 2F’ and data not shown).

The high percentage of sorting events (30% of all *ap* clones induced) raised the question of how many of these events were by chance, especially because the clones were induced prior to D/V boundary formation. To estimate the frequency of clones being born and twins ending in opposite compartments by chance, we analyzed control discs, where both sister clones were wild-type (Fig S1, 2G-I blue lines). In such control discs, nearly all dorsal clones (95%) remained in the dorsal compartment together with their twins (Fig 2G). The sister clones located in different compartments were observed very rarely (below 5%) (Fig 2H). Therefore, we conclude that dorsally originated *ap* mutant clones that are in close proximity to the D/V boundary are actively sorted into the ventral compartment.

Finally, we found that a high number of dorsal mutant clones (42%) were completely eliminated from the wing discs (Fig 2F, 2I). The majority of the elimination took place early, between the first two time points. Overall, 72% of all dorsal *ap* mutant clones were removed from the dorsal compartment: 30% *via* sorting and 42% *via* full elimination.

### The later the clone induction, the less efficient is the elimination

Next we asked whether the elimination and sorting rates depend on the clone induction time. To address this question, we induced the *ap* clones later, at 66h AEL (time after boundary formation), and analyzed at 100h and 110h AEL, which correspond to 34h and 44h after heat-shock (AHS), respectively (Fig 3A). Thus, we can compare the results of this experiment (late induced clones) with the results of our previous experiment (early induced clones) at least for 34 and 44h AHS.

**Fig 3.**
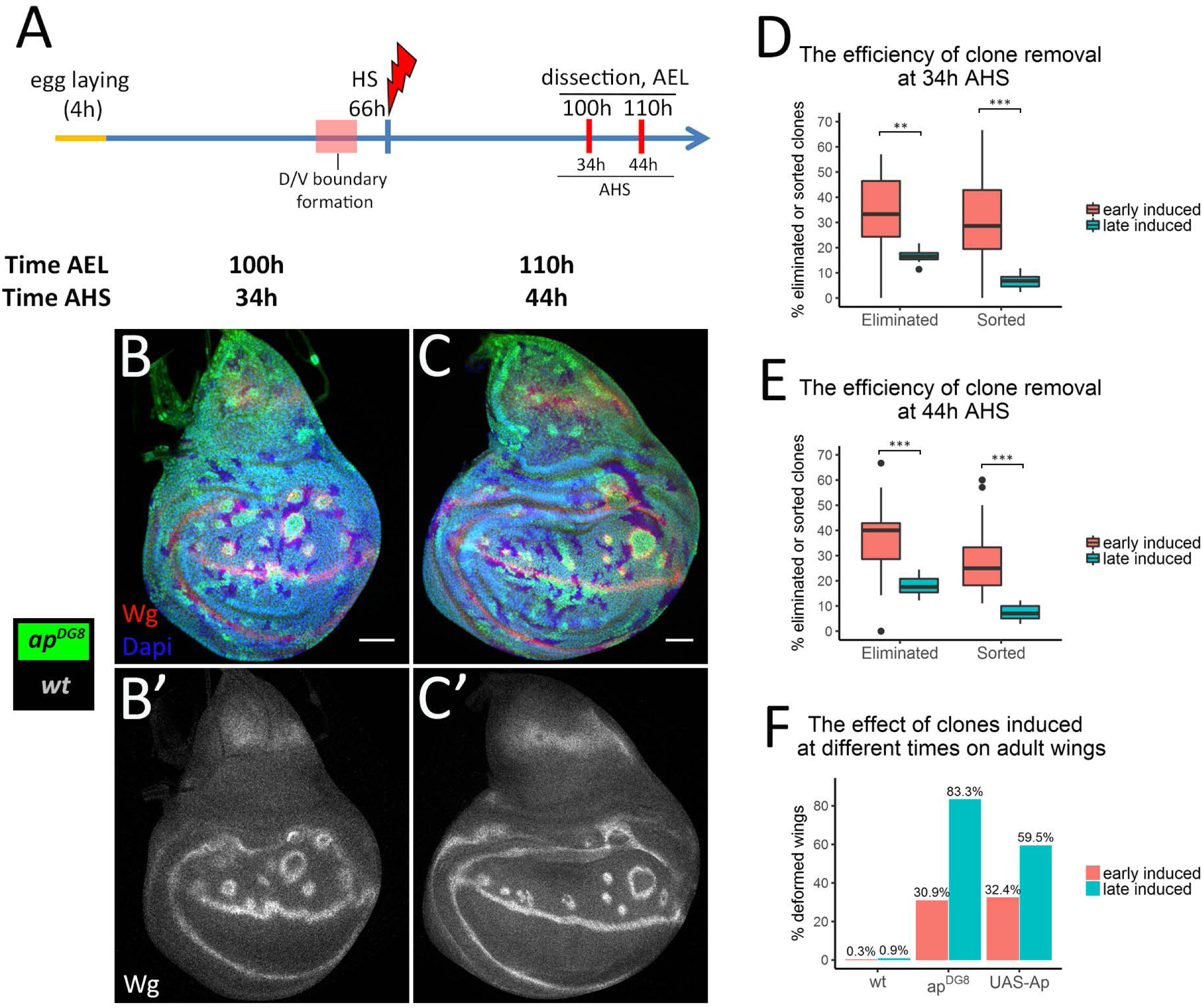
The late induced clones are eliminated less efficiently than the early induced ones. (A) Time-course scheme where clones are induced at 66h AEL. (B-C) Wing discs of indicated times containing differently marked *ap^DG8^* and wild-type clones. (B’-C’) Wg channel alone. (D-E) Comparison of removal of *ap^DG8^* clones that were induced at 46h AEL (“early induced”, shown in Fig2) with the one of those that were induced at 66h AEL (“late induced”) at 34h (D) and 44h (E) AHS. At least 10 discs with late induced clones were analysed per time point. (F) Percentages of defective wings due to wild-type, *ap^DG8^* and Ap-expressing flip-out clones induced at 46h or at 66h AEL. Numbers of analysed wings: early induced: wt - 302; *ap^DG8^* - 304; UAS-Ap – 482; late induced: wt - 230; *ap^DG8^* - 102; UAS-Ap – 84. Scale bars represent 50µm.

Expectedly, the clones were more abundant when induced in older discs due to the higher cell number (compare Fig 3B-C with Fig 2D-E). Using the categorization strategy described (Fig 2B), we quantified the percentage of mutant clones of dorsal origin that were either completely eliminated from the disc tissue or sorted to the ventral compartment. We found that the portions of completely eliminated mutant clones as well as sorted ones were significantly smaller for the late induced clones compared to the early induced ones at both 34h (Fig 3B, 3D) and 44h AHS (Fig 3C, 3E).

Remarkably, the induction of *ap* LOF clones after D/V boundary formation resulted in boundary deformations (Fig 3B’, note the wiggly D/V boundary) similar to what we observed when the clones were induced before the D/V boundary formation (Fig 2E’). Therefore, the compartment boundary can be rearranged after its formation. Such boundary flexibility allows the mutant clones to be rescued by displacement to the ventral compartment, though this happens more rarely for the clones induced after boundary formation than for the ones induced before. Altogether, the efficiency of misspecified cell elimination depends on the developmental stage: the mutant clones induced early are eliminated from the dorsal compartment more efficiently than the late induced ones.

Previous studies showed that *ap* mutant clones can lead to deformations in the adult wings (3). Indeed, in most cases, the presence of cells with inappropriate dorso-ventral positional identity in adult tissue caused ectopic margin formation (Fig S2B), wing margin duplications (Fig S2C, S2C’), blister-like outgrowths (Fig S2D, S2E), and, occasionally, wing duplications (Fig S2F, S2G). Importantly, the occurrence of defective wings highly correlates with the time of clone induction. When *ap* clones were induced late a vast majority of the wings (83%) had defects. In contrast, only one out of three wings were defective when the induction was early (Fig 3F). We observed the same trend with Ap-expressing clones induced at different times (Fig 3F). Thus, early induced misspecified clones are more likely to be eliminated, leading to normal wings. This finding highlights the importance of mechanisms in place to eliminate misspecified cells.

### Three region specific mechanisms ensure the clearance of misspecified cell clones

Next, we set out to study how the misspecified cells are being cleared from the tissue. As discussed, for *ap* mutant cells in close proximity of the D/V boundary, an elegant solution is to cross over to the opposite side (Fig 4A). This strategy also works for the clones misexpressing Ap (Fig 4B). During this process the ectopic boundary signaling induced between the misspecified cells and surrounding wild-type cells fuses with the regular D/V boundary, forming a loop-like structure around the misspecified clone (Fig 4A’, 4B’). This allows dorsal mutant cells or ventral Ap-expressing cells to mix with the cells from the opposite compartment and eventually recover at the correct place.

**Fig 4.**
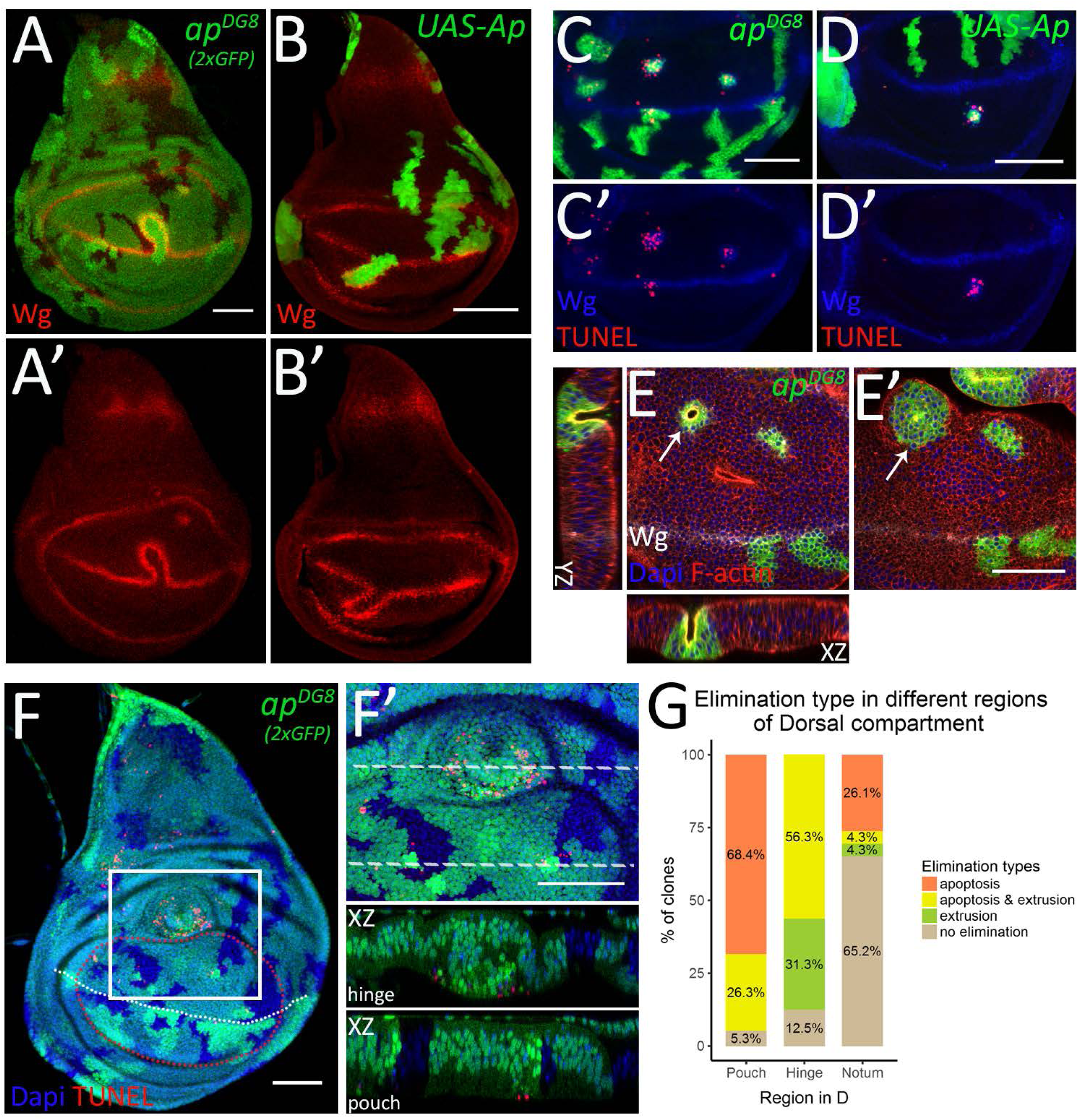
Mechanisms of the elimination display region specificity. (A-B) Third instar wing discs with *ap^DG8^* (A) or Ap-expressing (B) clones displaying D/V boundary deformation and clone sorting. (A’-B’) Wg channel of A-B. (C-D) TUNEL assay of third instar wing discs with *ap^DG8^* MARCM (C) or Ap-expressing (D) clones. The pouch regions are shown. Disc orientation: dorsal – up, ventral – down. (C’-D’) Wg and TUNEL channels of C-D. (E-E’) Pouch region of the third instar wing disc with *ap^DG8^* clones shown from the apical (E) and the basal (E’) sides. The arrows point to the bulging clone. The XZ and YZ planes throughout the bulging clone are also shown (XZ orientation: the apical side – up, YZ orientation: the apical side – right). (F-G) Mechanisms of elimination display region-specificity. (F) Third instar wing disc containing *ap^DG8^* mitotic clones (marked by 2 copies of GFP) that remained at 50h AHS. White dashed line represents D/V boundary; red dashed line outlines the pouch region. (F’) Zoom-in of the region defined by white square in F. The XZ cross-sections throughout the clones located in the hinge and the pouch are shown below (orientation: the apical side – up). (G) Quantification of dorsal mutant clones in different regions depending on the evidence of elimination type: apoptosis, extrusion or apoptosis together with extrusion. A total of 77 dorsal *ap* mutant clones from 23 discs were analyzed: 38 clones were in the pouch, 16 in the hinge and 23 in the notum. Scale bars represent 50µm.

Another mechanism contributing to the elimination of misspecified cells is apoptosis (11). Indeed, as revealed by TUNEL assay, both *ap* mutant and Ap-expressing clones undergo apoptosis in the inappropriate compartment (Fig 4C-D’). Interestingly, the apoptotic cells were detected both within and surrounding the misspecified cell clones (Fig 4C-D’).

Moreover, some misspecified clones displayed evidence of basal extrusion. The apical surfaces of those clones were narrower (Fig 4E), than their basal side (Fig 4E’), and the central cells were much shorter (Fig 4E, XZ and YZ). More lateral cells of the clone will eventually fuse above the gap forming a cyst-like structure with the apical side enclosed inside, contributing to the clearance.

Importantly, the vast majority of the bulging *ap* mutant clones were in the prospective hinge or in the very proximal pouch regions of the dorsal compartment. To estimate if there is any relationship between the region of clone location and the mechanism of elimination we carefully analyzed all *ap* mutant clones that remained in the dorsal compartment 50h after clone induction in the third instar wing discs. Moreover, the TUNEL assay allowed us to detect apoptotic cells. We reasoned that all sorting events at the boundary region had already taken place by this time. Thus, we focused only on the clones that were trapped in the dorsal compartment. Mutant clones from 23 wing discs were analyzed. Clones located in different regions of the dorsal compartment (dorsal pouch, dorsal hinge and the notum) were grouped based on the type of elimination they displayed: apoptosis (without evidence of extrusion), extrusion (without apoptosis), extrusion accompanied by apoptosis and the clones that did not display any evidence of elimination (Fig 4G). In the pouch, almost all misspecified clones underwent apoptosis (36 clones out of 38 clones examined in the pouch) (Fig 4F-G). Some of these apoptotic clones also bulged out, especially the ones in the proximal pouch (10 clones out of 36). We found no examples of extrusion for the clones located closer to the D/V boundary. In contrast, in the hinge, the majority of the mutant clones displayed a cyst-like phenotype (14 clones out of 16 examined clones in the hinge) (Fig 4F-G). Interestingly, the bulging clones in the hinge were not necessarily accompanied by apoptosis, but all apoptotic clones displayed extrusion (Fig 4G). This finding suggests that the induction of clone extrusion in the hinge is not a consequence of apoptosis. The opposite scenario is more likely - apoptosis takes place following clone extrusion in the hinge. In the notum, 7 clones out of 23 examined contained apoptotic cells, and only 2 clones formed invaginations (Fig 4G). However, the majority of remaining clones displayed evidence of neither apoptosis nor extrusion. Notably, the presence of the unmarked twin-spots in the central notum suggests that mutant clones had already been eliminated from this region by either apoptosis or extrusion.

Altogether these data indicate that misspecified cells are removed by three different mechanisms: sorting, apoptosis and basal extrusion. Moreover, apoptosis dominates in the pouch, whereas extrusion is the main mechanism in the hinge.

### Extrusion occurs independently of apoptosis and takes over in the absence of cell death

To assess the contribution of cell death to the elimination process, we prevented apoptosis in *ap^DG8^* cells by co-expression of the inhibitor of apoptosis *p35*. Wild-type, *UAS-p35*, *ap^DG8^* and *ap^DG8^* with *p35* clones were induced at early second instar and the discs of mid-third instar larvae were analyzed. The clones expressing only *p35* (Fig 5B) behaved similarly to the wild-type *GFP*-expressing clones (Fig 5A). In both cases, the clones did not display apoptosis, as revealed by the TUNEL assay. In contrast, dorsally located *ap* mutant clones induced apoptosis (Fig 5C). As expected, expression of *p35* within the clones perfectly inhibited apoptosis of clonal cells, but did not prevent induction of apoptosis outside the clone (Fig 5D, upper insert).The expression of *p35* in the mutant clones significantly increased the number of recovered clones (Fig 5D-E). However, the number of mutant clones expressing *p35* was still lower than that of wild-type or *p3*5-expressing clones (Fig 5E).

**Fig 5.**
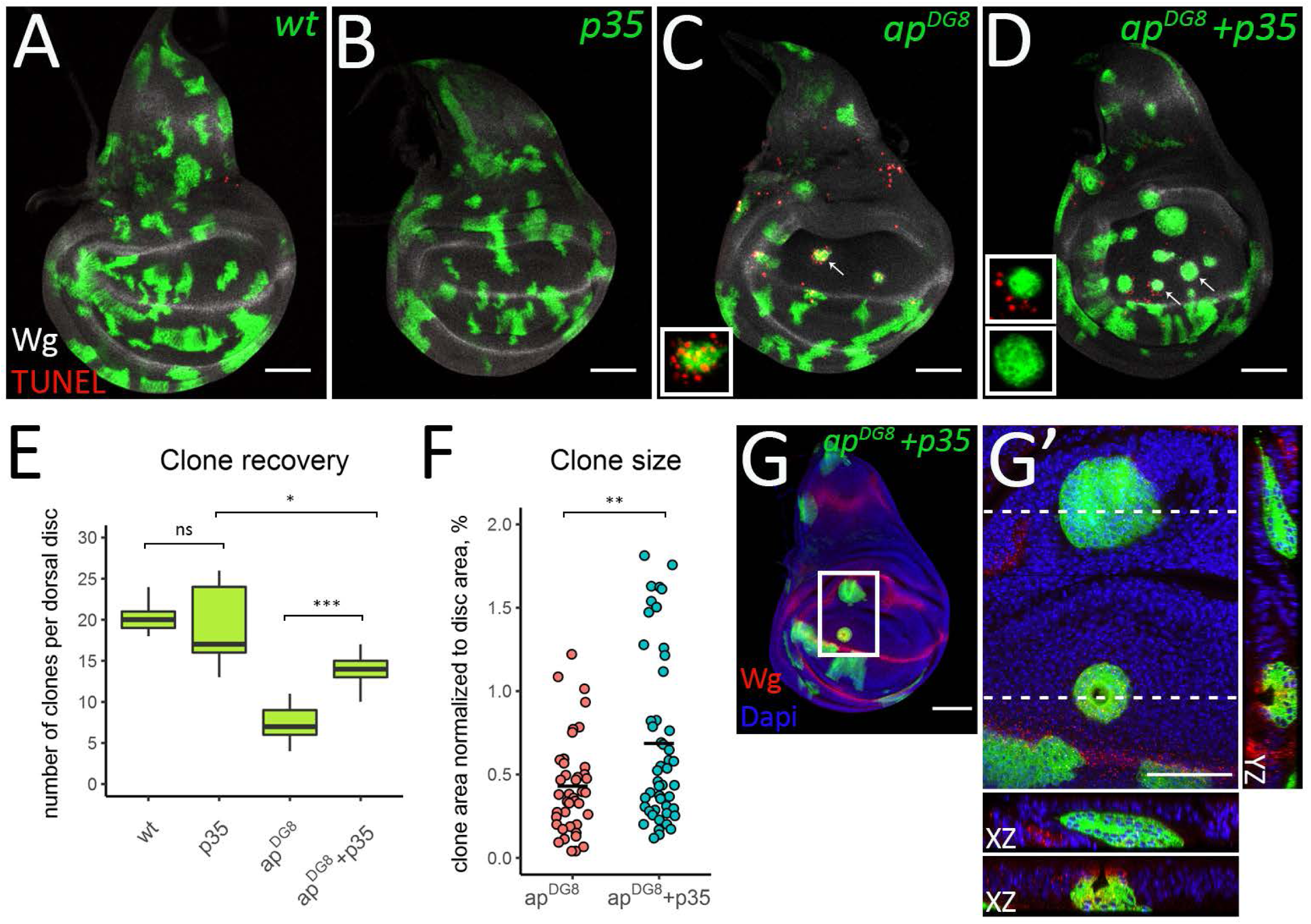
In the absence of apoptosis misspecified clones become bigger and undergo extrusion. (A-D) Third instar wing discs with wild-type (A), *p35*-expressing (B), *ap^DG8^* (C) and *ap^DG8^ p35*-expressing clones. The insets in C and D show enlarged images of single representative clones defined by arrows. (E) Clone recovery in the dorsal disc. 9 discs for each genotype were analyzed. (F) Plot shows areas of *ap^DG8^* and *ap^DG8^ p35*-expressing clones. (G-G’) *ap* mutant clones are eliminated from the dorsal pouch via extrusion when apoptosis is blocked. (G) Third instar wing disc containing *ap^DG8^ p35-*expressing clones. (G’) Zoom-in of the region defined by the white square in G. The XZ and YZ cross-sections throughout the clones located in the hinge and in the pouch are shown (XZ orientation: the apical side – up; YZ orientation: the apical side - left). Scale bars represent 50µm.

Thus, apoptosis inhibition only partially rescues the elimination of the misspecified clones. To determine whether apoptosis inhibition influenced the sorting efficiency and whether the clones are indeed eliminated from the tissue in the absence of apoptosis, we analyzed *ap^DG8^* clones expressing *p35* together with their wild-type sisters. The clones were induced at 46h AEL and analyzed at 80h, 90h and 100h AEL (similar to our time-course experiment in Fig 2). The quantification of sorting events and comparison to the results obtained with the *ap* mutant clones alone (Fig 2), revealed that the expression of *p35* did not change the sorting efficiency at any time-point (Fig S3A-D). Approximately 30% of clones were sorted to the ventral compartment (Fig S3D). In contrast, the number of mutant clones that were fully eliminated from the disc tissue were significantly lower when apoptosis was blocked (Fig S3E). However, about 18% of dorsal wild-type clones were found without their mutant sisters (Fig 3SB-C, arrows and 3SE). This directly indicates that the misspecified clones can be eliminated from the tissue even in the absence of apoptosis. Many *ap* mutant clones with *p35* expression displayed evidence of basal extrusion. Interestingly, in this case cyst formation was observed not only in the hinge region but also in the notum and in the pouch (Fig 5G-G’). This suggests that extrusion does not depend on apoptosis and can serve as a back-up mechanism of clone elimination.

### Clone size is important for cyst-formation but not for clone elimination via extrusion

One possible explanation of why the hinge clones but not the pouch ones undergo extrusion is the clone size. Misspecified clones in the hinge are larger than the ones in the pouch (Fig 4F). Therefore, we wondered whether clone size would be linked to the choice of elimination mechanism. This could also explain why clones in the pouch (and in the notum) begin extruding upon apoptosis inhibition: since many *ap* mutant clones in the pouch normally undergo apoptosis, they may not have a chance to reach the size required for extrusion, while apoptosis inhibition allows mutant clones to grow and reach a larger size (Fig 5F).

Thus, we asked whether changing the clone size affects their elimination in the presence and absence of apoptosis. To reduce the clone size we made use of *string (stg)* RNAi. Stg is an activator of the cyclin-dependent kinases. It regulates cell cycle progression by driving cells into mitosis (35). Accordingly, *stgRNAi* expressing cells proliferate slowly and the clones have smaller size compared to the wild-type clones (Fig S4A, S4C). However and importantly the expression of *stgRNAi* did not affect the clone recovery rate (Fig S4I). Therefore, *stgRNAi* expression and associated with it reduction of proliferation do not cause clone elimination per se. As previously, *p35* was used to prevent apoptosis within the clones (Fig S4B). The clones expressing both *p35* and *stgRNAi* combined both effects: they were smaller, and were recovered at a higher rate than wild-type clones (Fig S4D, S4I). To modulate apoptosis and clone size in misspecified cells at the same time, we used *dLMO* flip-out clones instead of *ap* mitotic clones. Like *ap* mutant clones, *dLMO* flip-out clones induce ectopic boundary signaling and are efficiently eliminated from the dorsal compartment (Fig 6A, S4E, S4I; see also (11, 36)). The distribution of the elimination types the *dLMO* clones were undergoing in different regions mimics the one of *ap* mutant clones (Fig 6E, *dLMO*). Upon apoptosis inhibition the behavior of *dLMO* clones again resembled the behavior of *ap* mutant clones (Fig 6B, S4F): *p35* co-expression increased the clone recovery rate (Fig S4I) and led to clone extrusion in all regions of dorsal compartment (Fig 6E, *dLMO + p35*). Contrary to our expectations, the reduction of *dLMO* clone size did not influence the clone recovery rate (Fig 6C, S4G, S4I). These smaller *dLMO* clones were less likely to be associated with apoptosis and more frequently displayed invagination, shortening, and extrusion phenotypes, especially in the pouch and in the notum (Fig 6E, *dLMO + stgRNAi*). Moreover, when we reduce the size of *dLMO* expressing clones and prevent their apoptosis at the same time, most of misspecified clones underwent extrusion in all parts of the dorsal compartment (Fig 6D, 6E, *dLMO + stgRNAi + p35*). The fact that reduction of clone size does not prevent clone extrusion and does not increase the clone recovery rate suggest that elimination of misspecified clones via extrusion occurs regardless of the clone size.

**Fig 6.**
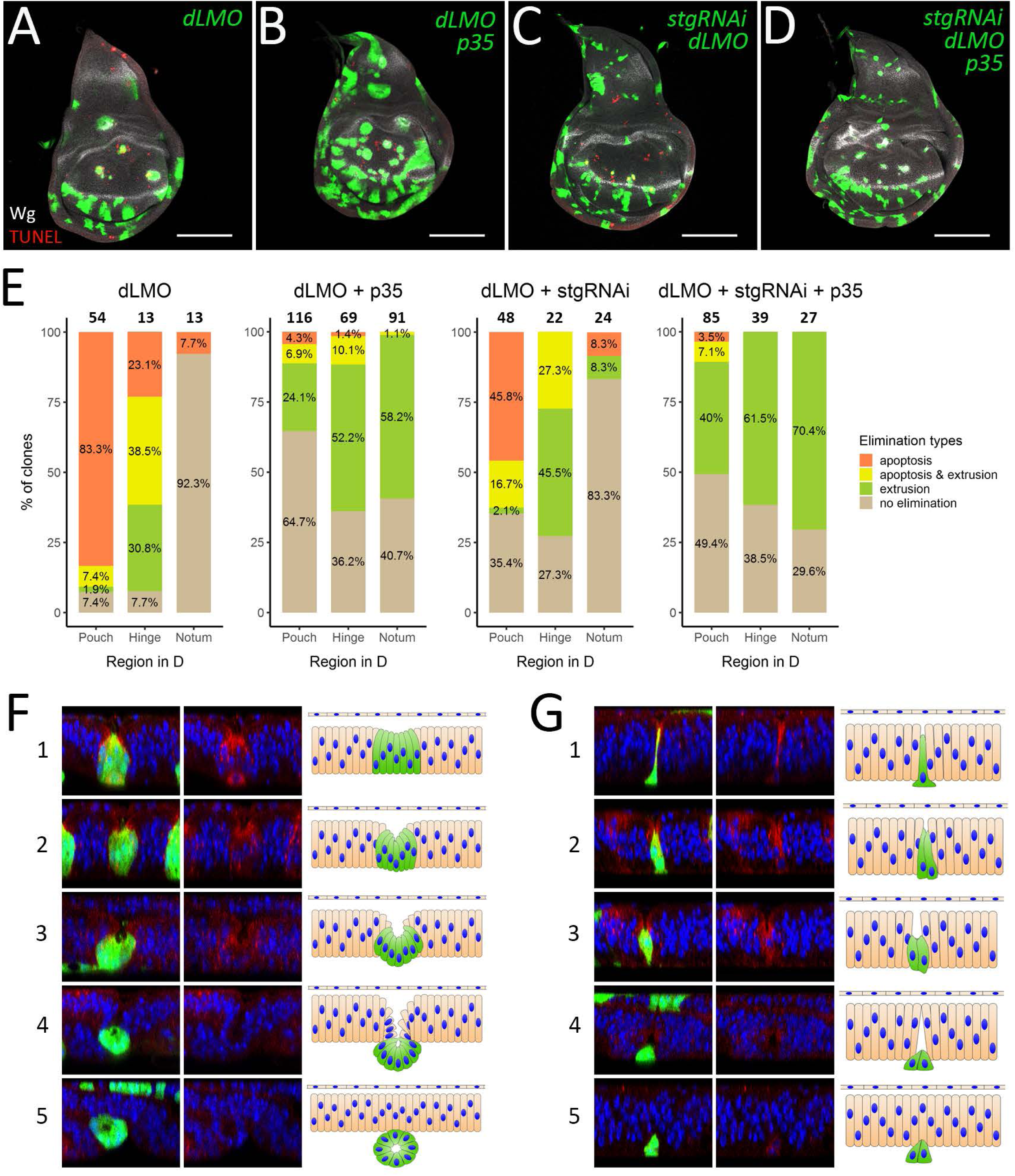
Clone size reduction does not prevent clone extrusion. (A-D) Third instar wing discs containing *dLMO* (A), *dLMO+p35* (B), *stgRNAi+dLMO* (C) and *stgRNAi+dLMO+p35* (D) clones. (E) Quantification of clones of the indicated genotypes in different regions of dorsal compartment depending on the evidence of elimination type: apoptosis, extrusion or apoptosis together with extrusion. The numbers of analyzed clones in each region are displayed above the bars. (F-G) Extrusion of large and small clones. Examples of large (*dLMO+p35*) (F) and small (*stgRNAi+dLMO+p35*) (G) misspecified clones at different stages (1–5) of extrusion process. XZ cross-sections throughout the clones are shown (orientation: the apical side – up). The left panel: clones in green, nuclear staining (Dapi) in blue, Wg staining in red; the middle panel - nuclear and Wg staining alone; right panel – schematic representation of morphological changes. The initial steps of extrusion of both large and small clones involve apical constriction and reduction of cell height (cell shortening) from the apical side (F and G, 1-2). Propagation of those processes, especially cell shortening, in case of large clones leads to cyst formation, where apical sides of clonal cells face the newly-formed cavity (F, 3-4). In contrast, small clones do not form cysts, although the cells reduce their height further and get extruded from the tissue (G, 3-4). Finally, the wild-type neighboring cells fuse above the clones and restore epithelium integrity (F and G, 5). Scale bars represent 100µm.

Although the small size does not prevent the misspecified clones from being extruded, such clones extrude from the tissue in a different way than the larger ones do. Careful analysis of *dLMO + p35* and *dLMO + p35 + stgRNAi* clone morphology showed that *dLMO + p35* clones, which were generally of medium to large size (more than 6 cells), form cyst-like structures with a cavity inside, whereas small clones (1 - 6 cells) did not. Most clones expressing *stgRNAi* were clones of the small size. During our analysis, we found clones at different steps of extrusion process from which we could reconstruct the whole process for both the large (Fig 6F) and the small (Fig 6G) clones. At the first step the large clones experience shrinkage of the apical surface (Fig 6F-1). At the same time clonal cells, especially cells in the clone center, get shorter leading to clone invagination (Fig 6F-2). Eventually all cells in the clone are reduced in height and the clone forms a cyst-like structure (Fig 6F-3). The cyst is pushed out from the tissue plane and becomes enclosed (Fig 6F-4). After the cyst extrusion is complete, the disc epithelium restores its integrity (Fig 6F-5). The small clones also begin the extrusion process by constriction of their apical areas, expansion of the basal side and cell shortening (Fig 6G-1). These changes lead to local tissue invagination (Fig 6G-2, -3). Further reduction of clone height causes clone extrusion. At the same time, neighboring wild-type cells establish contacts above the extruding clone (Fig 6G-4) and the tissue restores its integrity and shape (Fig 6G-5). In conclusion, unlike large clones, small clones do not form cyst-like structures, however both types of clones can leave the tissue via basal extrusion. The apical construction and cell shortening are the common changes resulting in local tissue invagination and clone extrusion.

## DISCUSSION

Here we studied the behavior of cells misspecified for the dorso-ventral identity. Using a non-canonical FRT site (33) we induced *ap* mutant clones and analyzed their behavior in the dorsal compartment. Interestingly, the misspecified cells are not eliminated immediately after induction. Initially, we suspected that the clones needed to reach a certain size to initiate elimination. However, our data shows that the clone size is not a decisive parameter for the elimination. The misspecified cell clones, as small as 1 cell, can be extruded from the epithelia. Alternatively, the effect might be due to Ap protein or the transcript stability. In this scenario, bringing Ap below a certain threshold simply requires time or several divisions that would dilute the protein level in each cell. Although the misspecified clones are able to grow within the first 24h, most of them are recognized and effectively eliminated from the dorsal compartment within the following 10h. We have defined 3 mechanisms that ensure their elimination: sorting to the opposite compartment, apoptosis and basal extrusion.

### Sorting to the opposite compartment

The phenomenon, when dorsal cells mutant for *ap* cross the boundary and join the ventral compartment, has been observed in the early work defining Ap as the dorsal determinant (32). Importantly, this ability of cells to swap compartments according to their identity contributes to the elimination of misspecified cells. We found that up to 30% of mutant clones of dorsal origin leave the compartment via this mechanism. Three main events make clone sorting possible: the induction of ectopic boundary signaling around the clone, incorporation of this signaling into the compartment boundary, leading to loops protruding from the D/V boundary, and boundary straightening. How boundary straightening occurs is not known. However, it is very likely that the mechanisms that maintain the boundary straight during normal development are in effect here. For instance, it was shown that the D/V boundary has distinct physical parameters such as increased cell bond tension, cell elongation and oriented cell division, which tightly correlate with the boundary morphology and ensure its straightness (37). Importantly, the increased tension depends on Ap and Notch activity (38). Therefore, it is possible that the mechanical changes associated with displaced signaling help to bring D/V boundary to the normal shape. Notably, the ability of misspecified dorsal clones cross into the ventral compartment even after D/V boundary formation suggests that the signaling center is a very flexible and dynamic structure. It can be rearranged at any time during development in order to meet tissue needs.

### Apoptosis and Extrusion

Although some misspecified clones, the ones that are close to the D/V boundary, can escape to the opposite compartment and survive, the majority of misspecified clones are completely eliminated from the disc tissue either via apoptosis or basal extrusion. Interestingly, apoptosis activation occurs in both the misspecified cells and the juxtaposing wild type cells. Moreover, inhibition of apoptosis in the clones does not prevent its non-autonomous activation. This suggests that apoptosis activation rely rather on interaction of cells with different fate identities than on misspecified cells themselves. Similar autonomous and non-autonomous activation of apoptosis was reported for the adjacent cells that experience discontinuity in the reception of either the Dpp or Wg signaling (21).

A former study by Marco Milan and colleagues reported that *p35* co-expression rescued dLMO clones of dorsal origin completely (11). In our set up the rescue effect of *p35* was also significant, however incomplete (Fig 5E and S3). We find that in addition to apoptosis, basal extrusion also contributes to the elimination of cells misspecified for the D/V position. The underlying reason of the discrepancy between the published results and ours might be the timing of clone induction, as the later induced clones are more likely to escape the elimination mechanisms in place. Another important factor that could contribute to the differences between *p35* rescue experiments is that the analysis in Milan paper was restricted to the pouch region, whereas we analyzed clones throughout the whole dorsal compartment. Therefore, we suspect that earlier clone induction along with quantification in the whole disc allowed us to recognize the contribution of basal extrusion to the process of clone elimination.

### Region specificity

Interestingly, apoptosis and extrusion display strong region preferences: apoptosis dominates in the pouch whereas extrusion occurs preferentially in the hinge. Such an interesting pattern could rely on three factors. First, the cyto-architectural properties of the hinge and the pouch regions are different. Cells in the wing pouch have a long and narrow shape along their apical-basal axes, whereas cells in the hinge are shorter and wider (13, 39). This makes the hinge region mechanically more disposed to bulging (40). Second, the hinge region is resistant to irradiation and drug-induced apoptosis due to low levels of the pro-apoptotic gene *reaper* in that region (41). Third, we find that *ap* mutant clones in the dorsal pouch and dorsal hinge have different effects on cell proliferation. The misspecified clones in the hinge increase cell proliferation in both an autonomous and a non-autonomous manner. By contrast, the clones in the pouch either grow at the normal rate or even slightly inhibit cell proliferation (Fig S5). Most likely these effects are mediated by ectopic Notch/Wg signaling induced at the clone boundary. Indeed, it was reported that Notch or Wg misexpression increases cell proliferation and causes strong overgrowth in the hinge but not in the pouch (10, 13, 23, 42). Thus, the extrusion of misspecified clones in the hinge could be driven by local crowding, which was shown to be linked to extrusion in the *Drosophila* pupal notum (43, 44). However, our data suggests that the role of local crowding can be at most minor with regards to the extrusion of *ap* mutant clones. First of all, in the absence of apoptosis, the extrusion of *ap* clones occurs rather frequently not only in the hinge but also in the pouch and in the notum, where the clones do not induce overgrowth. In addition, the clones with artificially reduced size (*dLMO+p35+stgRNAi*) are still extruded from the epithelium despite the lack of the crowding effect (Fig 6E and 6G). Overall, we think that it is the in-built apoptotic resistance in the hinge, rather than differential proliferation patterns that favors extrusion in the hinge. Here, we described three mechanisms that ensure clearance of cells with incorrect D/V identity and their regional bias. We also find that the elimination of misspecified cells is more efficient earlier in development. This suggests that the ability of developing tissue to remove inappropriately specified cells and actively maintain the compartment organization requires some tissue plasticity that diminishes over time.

## MATERIALS AND METHODS

### Fly stocks

The following fly stocks were used in this study: *ap^DG8^* (described in Bieli et al., 2015), *FRT^f00878^* (described in Bieli et al., 2015), *UAS-dLMO* (was kindly provided by Marco Milan), *UAS-Ap* (was kindly provided by Markus Affolter), *UAS-p35* (was kindly provided by Nicole Grieder); *UAS-stgRNAi* (GD, 330033) obtained from the Vienna Drosophila Resource Center (VDRC). All crosses were kept on standard media at 25*^◦^*C. Flipase expression was induced by a heat-shock at 37*^◦^*C. The detailed fly genotypes and heat-shock induction conditions are presented in Table S1.

### Immunostaining and sample preparation

Imaginal discs were prepared and stained using standard procedures. Briefly, larvae were dissected and fixed in 4% paraformaldehyde (PFA) in PBS for 20 min. Washes were performed in PBS + 0.03% Triton X-100 (PBT) and blocking in PBT+2% normal donkey serum (PBTN). Samples were incubated with primary antibodies overnight at 4*^◦^*C. The primary antibodies used: mouse anti-Wg (1:2000, deposited to the DSHB by Cohen, S.M. (DSHB Hybridoma Product 4D4-concentrated)). Secondary antibodies were incubated for 2hr at room temperature. The secondary antibodies used: anti-mouse Alexa 568 and Alexa 633. Discs were mounted in Vectashield antifade mounting medium with Dapi (Vector Laboratories). For F-actin staining Phalloidin-Tetramethylrhodamine B (Fluka #77418) was added during incubation with secondary antibodies at the concentration 0.3 µM. For adult wing sample preparation, the flies of desired genotypes were collected and fixed in 70% ethanol. The wings were isolated and mounted in 3:1 Canadian balsam : Methyl Salicylate.

### TUNEL assay

For the TUNEL assay In Situ Cell Death Detection kit, TMR red (Roche) was used. Larvae were dissected in cold PBS and fixed in 4% PFA for 1hr at 4*^◦^*C. Samples were washed in PBT and blocked in PBTN for 1 hr. Next, samples were incubated with primary antibodies overnight at 4*^◦^*C and with secondary antibodies for 4hr at 4*^◦^*C. After washing the tissues were blocked in PBTN overnight at 4*^◦^*C. Then, samples were permeabilized in 100 mM sodium citrate supplemented with 0.1% Triton X-100 and incubated in 50 µl of TUNEL reaction mix (prepared according to the recipe from the kit) for 2 hr at 37*^◦^*C in dark. After this step, the samples were washed in PBT for 30 min and mounted in Vectashield antifade mounting medium with Dapi (Vector Laboratories).

### EdU labeling

For the EdU assay Click-iT EdU Alexa Fluor 594 imaging kit (Invitrogen #C10339) was used. Larvae were dissected in Schneider’s Medium at room temperature and incubated for 1 hr at 25°C in 15 µM EdU working solution supplemented with 1% normal donkey serum. After the EdU incorporation, the tissue was washed in PBS supplemented with 3% bovine serum albumin (BSA) and fixed in 4% PFA for 20 min. Next steps, including blocking, incubation with primary and secondary antibodies, were done according to the standard immunostaining protocol. After washing the tissues were permeabilized by 3 washes (10 minutes each) in 0.1% Triton X-100 in PBS. The EdU reaction cocktail was prepared according to the recipe from the kit. The samples were incubated in 250 µl of the EdU reaction cocktail for 30 min at room temperature in dark. After that the samples were washed in PBT for 30 min and mounted in Vectashield antifade mounting medium with Dapi (Vector Laboratories).

### Image acquisition and analysis

Image stacks of wing discs were acquired on Zeiss LSM710 or LSM880 confocal microscopes using 20x or 40x objectives. In most cases 15-30 Z-sections 1 µm apart were collected. Image stacks were projected using maximum projection and analyzed using workflows established in ImageJ. The images shown on Fig 4E-E’, 4F’, 5G’, 6F, 6G and all images used for the analysis of the elimination type (Fig 4G and 6E) were acquired using a 40x objective. In this case, 80-130 Z-sections 0.4-0.7 µm apart were collected. The orthogonal views throughout clone centers were used to define clones under extrusion. Statistical analysis was done in R, v3.5.0. Conditions were compared using two-sample t-test. Comparisons with a p-value > 0.05 were marked as “ns” (non-significant); p-value ≤ 0.05 – “*”; p-value ≤ 0.01 – “**”; p-value ≤ 0.001 – “***”.

## ACKNOWLEDGEMENTS

We are grateful to M. Müller, D. Bieli and M. Affolter of Biozentrum Basel for reagents and discussions. We thank M. Milan (IRB, Barcelona) and VDRC for providing fly stocks and DSHB for anti-Wg antibodies. Confocal microscopy was performed at the Cellular Imaging Facility of the University of Lausanne. We thank V. Dion for comments on the manuscript.

## SUPPORTING INFORMATION CAPTIONS

**Fig S1.**
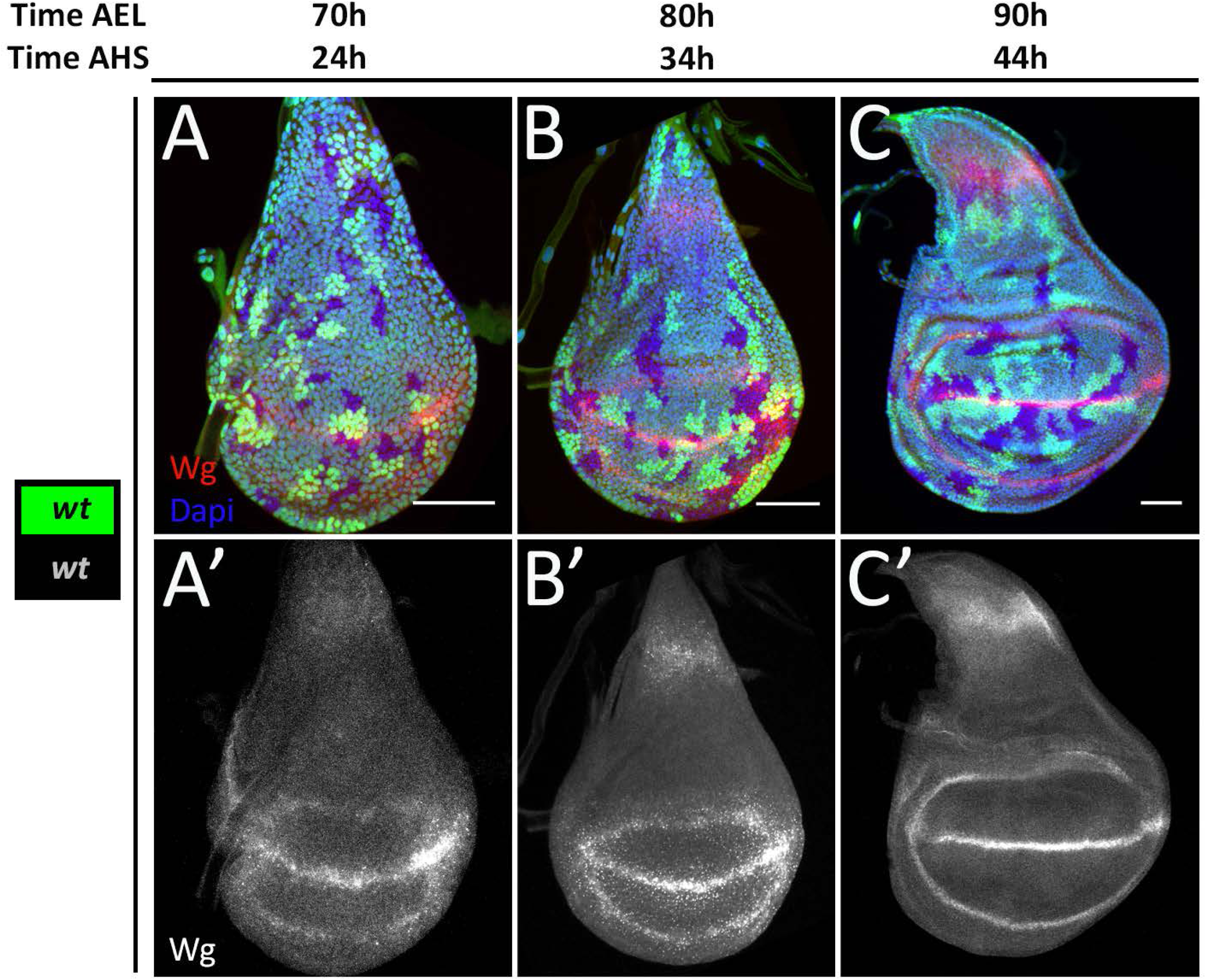
Sorting and elimination of wild type clones in the control discs are very rare events. (A-C) Wing discs of the indicated times containing wild-type sister clones that are marked by either 2 copies of GFP or absence of GFP. (A’-C’) Wg channel of A-C. Quantifications of the remained, sorted and eliminated wild-type clones are shown on the Fig 2G-I, blue lines. Scale bars represent 50µm.

**Fig S2.**
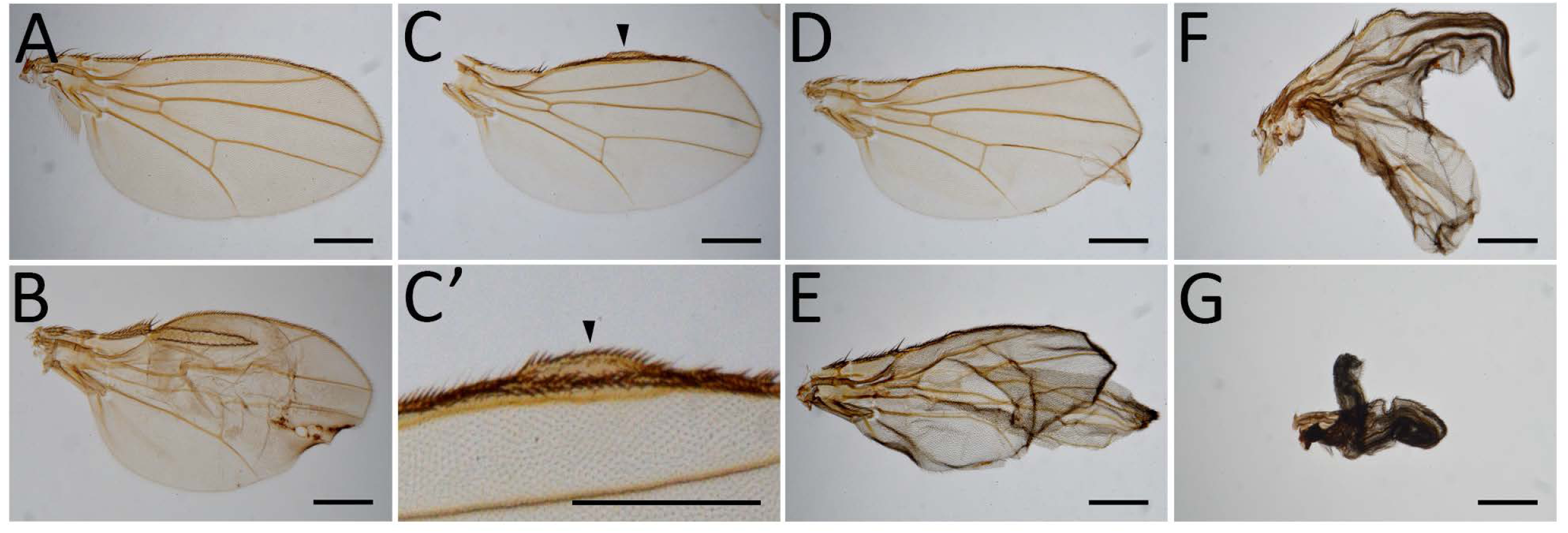
*ap* mutant cell clones cause deformations in adult wings. (A) Wild-type wing. (B-G) Wings after induction of *ap^DG8^* clones during second instar contain different deformations: ectopic margin formation (B); wing margin duplication (C-C’, arrowheads); blister-like outgrowths (D-E); and wing duplications (F-G). Scale bars represent 500µm.

**Fig S3.**
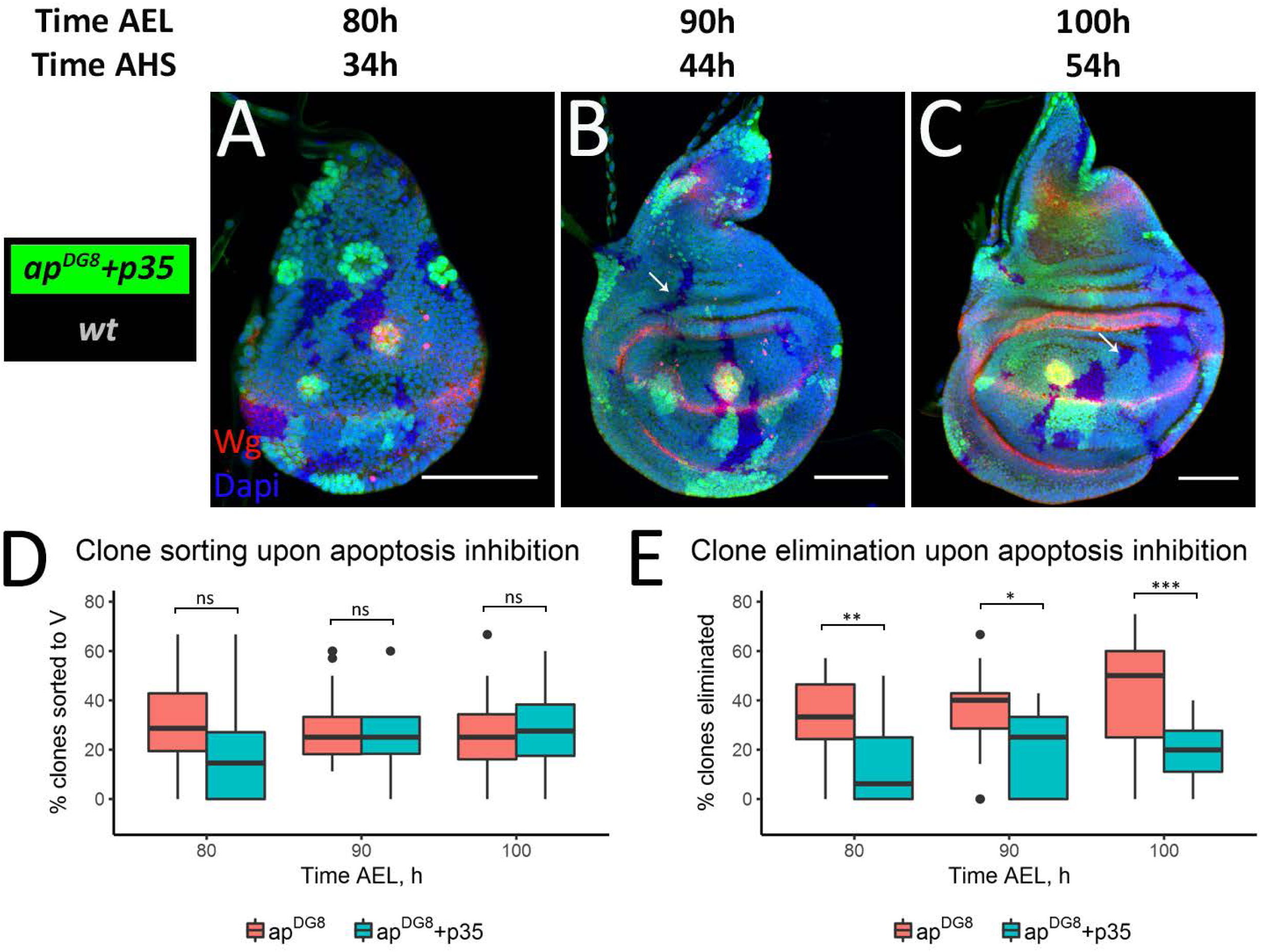
Apoptosis inhibition does not rescue all mis-specified clones. (A-C) Wing imaginal discs of indicated times with *ap^DG8^* clones expressing *p35* (marked by two copies of GFP) and wild-type sister clones (marked by the absence of GFP). Arrows point to wild-type clones that lost their mutant sisters; (D-E) Comparison of the amount of *ap^DG8^* clones (data from the Fig 2) with the amount of *ap^DG8^* + *p35* clones that were sorted to the ventral compartment (D) or completely eliminated (E). At least 15 discs with *ap^DG8^* clones and 12 discs with *ap^DG8^* + *p35* clones were analyzed. Scale bars represent 50µm.

**Fig S4.**
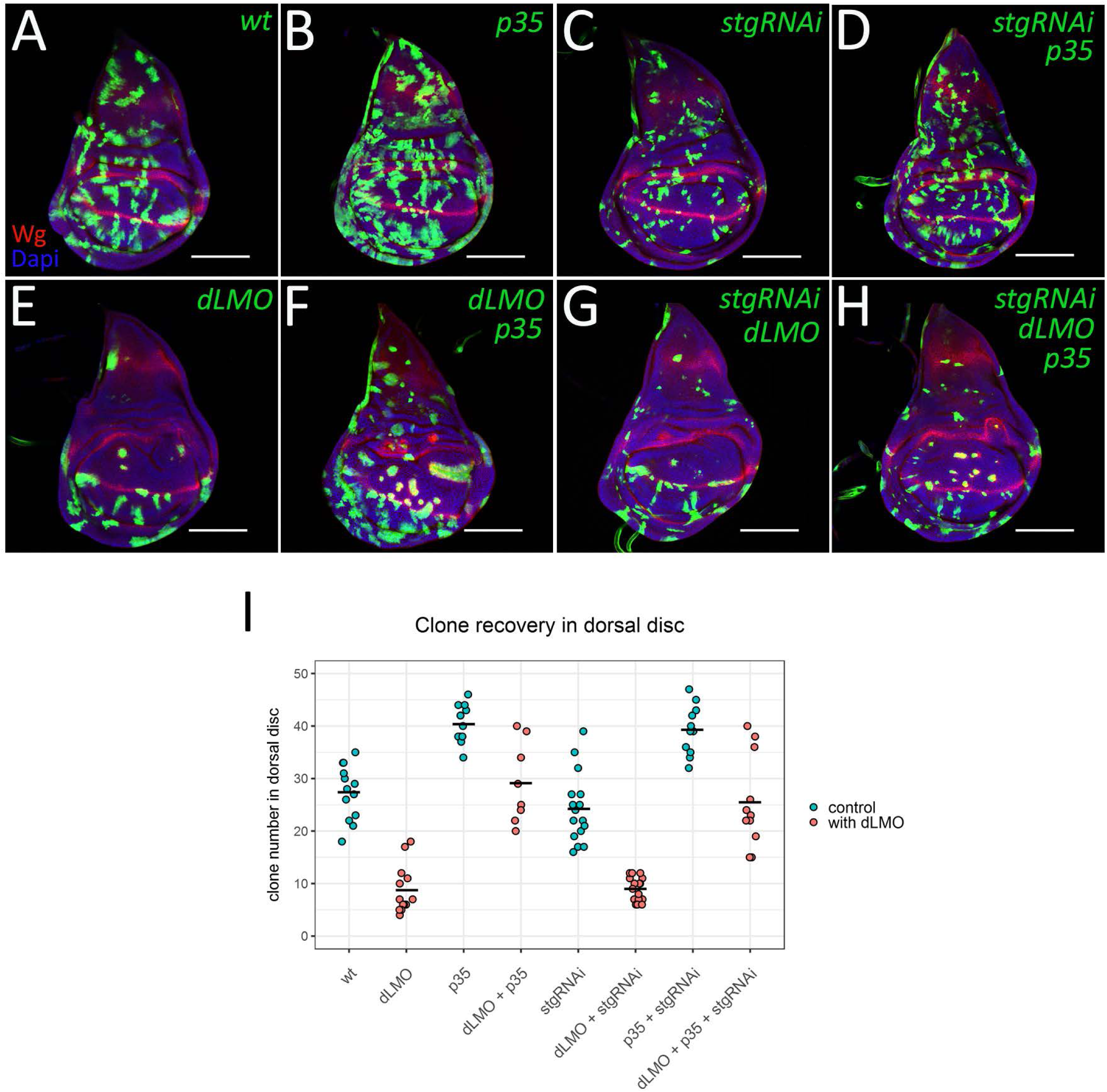
The reduction of clone size does not affect their recovery. (A-H) Third instar wing discs containing wild-type (A), *p35* (B), *stgRNAi* (C), p35+stgRNAi (D), *dLMO* (E), *dLMO+p35* (F), *stgRNAi+dLMO* (G) and *stgRNAi+dLMO+p35* (H) clones. (I) Clone recovery rate in dorsal compartment for each genotype. Scale bars represent 100µm.

**Fig S5.**
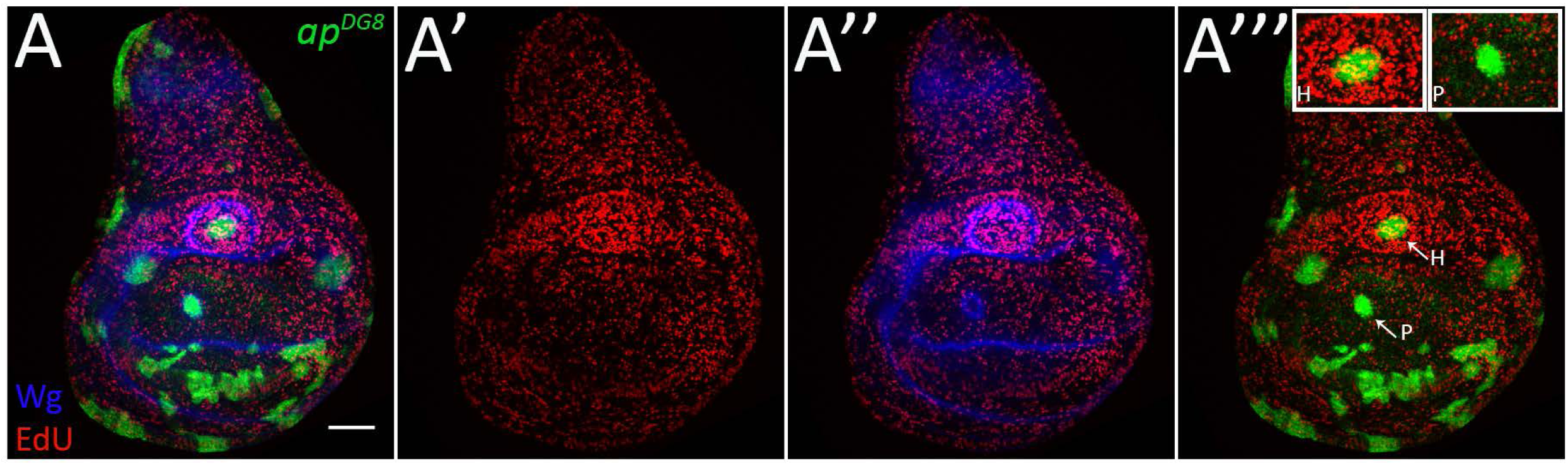
*ap* mutant clones increase cell proliferation in the dorsal hinge but not in the dorsal pouch. EdU cell proliferation assay of the third instar wing disc containing *ap^DG8^* clones. (A) Merged image (*ap^DG8^* clones, EdU and Wg staining). (A’) EdU channel alone. (A’’) EdU and Wg channels. (A’’’) *ap^DG8^* clones and EdU staining. The insets show enlarge images of single clones from dorsal pouch (P) and dorsal hinge (H). Scale bar represents 50µm.

**Table S1.** Genotypes and experimental conditions. Detailed genotypes and experimental conditions of data represented on individual figure.

| Figure | Genotype | Time AEL, h |  |  | Heat-shock duration, min |
| --- | --- | --- | --- | --- | --- |
|  |  | Egg collection | Heat-shock | Dissection |  |
| 1A | *yw hsflp /yw; FRT<sup>f00878</sup> / FRT<sup>f00878</sup> tub-Gal80; tub-Gal4 UAS-GFP/+* | 4 | 48-52 | 110-114 | 30 |
| 1B | *yw hsflp /yw; FRT<sup>f00878</sup> ap<sup>DG8</sup> / FRT<sup>f00878</sup> tub-Gal80; tub-Gal4 UAS-GFP / +* | 4 | 48-52 | 110-114 | 30 |
| 1C | *yw hsflp / w; UAS-Ap / +; act>CD2>Gal4 UAS-GFP / +* | 4 | 46-50 | 100-104 | 12 |
| 2 B | *yw hsflp/(y)w; FRT<sup>f00878</sup> ap<sup>DG8</sup> ubi-GFP / FRT<sup>f00878</sup>* | 4 | 42-46 | 86-90 | 30 |
| 2C | *yw hsflp/(y)w; FRT<sup>f00878</sup> ap<sup>DG8</sup> ubi-GFP / FRT<sup>f00878</sup>* | 4 | 42-46 | 66-70 | 30 |
| 2D | *yw hsflp/(y)w; FRT<sup>f00878</sup> ap<sup>DG8</sup> ubi-GFP / FRT<sup>f00878</sup>* | 4 | 42-46 | 76-80 | 30 |
| 2E | *yw hsflp/(y)w; FRT<sup>f00878</sup> ap<sup>DG8</sup> ubi-GFP / FRT<sup>f00878</sup>* | 4 | 42-46 | 86-90 | 30 |
| 2F | *yw hsflp/(y)w; FRT<sup>f00878</sup> ap<sup>DG8</sup> ubi-GFP / FRT<sup>f00878</sup>* | 4 | 42-46 | 96-100 | 30 |
| 3B | *yw hsflp/(y)w; FRT<sup>f00878</sup> ap<sup>DG8</sup> ubi-GFP / FRT<sup>f00878</sup>* | 4 | 62-66 | 96-100 | 30 |
| 3C | *yw hsflp/(y)w; FRT<sup>f00878</sup> ap<sup>DG8</sup> ubi-GFP / FRT<sup>f00878</sup>* | 4 | 62-66 | 106-110 | 30 |
| 4A | *yw hsflp/(y)w; FRT<sup>f00878</sup> ap<sup>DG8</sup> ubi-GFP / FRT<sup>f00878</sup>* | 4 | 42-46 | 86-90 | 30 |
| 4B | *yw hsflp / w; UAS-Ap / +; act>CD2>Gal4 UAS-GFP / +* | 4 | 42-46 | 86-90 | 12 |
| 4C | *yw hsflp /yw; FRT<sup>f00878</sup> ap<sup>DG8</sup> / FRT<sup>f00878</sup> tub-Gal80; tub-Gal4 UAS-GFP / +* | 24 | 32-56 | 76-100 | 30 |
| 4D | *yw hsflp / w; UAS-Ap / +; act>CD2>Gal4 UAS-GFP / +* | 4 | 42-46 | 86-90 | 12 |
| 4E | *yw hsflp /yw; FRT<sup>f00878</sup> ap<sup>DG8</sup> / FRT<sup>f00878</sup> tub-Gal80; tub-Gal4 UAS-GFP / +* | 8 | 51-59 | 92-100 | 30 |
| 4F | *yw hsflp/(y)w; FRT<sup>f00878</sup> ap<sup>DG8</sup> ubi-GFP / FRT<sup>f00878</sup>* | 8 | 40-48 | 92-100 | 30 |
| 5A | *yw hsflp /yw; FRT<sup>f00878</sup> / FRT<sup>f00878</sup> tub-Gal80; tub-Gal4 UAS-GFP/+* | 24 | 32-56 | 76-100 | 30 |
| 5B | *yw hsflp /yw; FRT<sup>f00878</sup> / FRT<sup>f00878</sup> tub-Gal80; tub-Gal4 UAS-GFP/ UAS-p35* | 24 | 32-56 | 76-100 | 30 |
| 5C | *yw hsflp /yw; FRT<sup>f00878</sup> ap<sup>DG8</sup> / FRT<sup>f00878</sup> tub-Gal80; tub-Gal4 UAS-GFP/+* | 24 | 32-56 | 76-100 | 30 |
| 5D | *yw hsflp /yw; FRT<sup>f00878</sup> ap<sup>DG8</sup> / FRT<sup>f00878</sup> tub-Gal80; tub-Gal4 UAS-GFP/ UAS-p35* | 24 | 32-56 | 76-100 | 30 |
| 5G | *yw hsflp /yw; FRT<sup>f00878</sup> ap<sup>DG8</sup> / FRT<sup>f00878</sup> tub-Gal80; tub-Gal4 UAS-GFP/ UAS-p35* | 24 | 24-48 | 96-120 | 30 |
| 6A | *yw hsflp / w; UAS-dLMO / IF or CyO; act>CD2>Gal4 UAS-GFP / MKRS* | 8 | 60-68 | 106-114 | 13 |
| 6B | *yw hsflp / hsflp; UAS-dLMO / CyO or IF; act>CD2>Gal4 UAS-GFP / UAS-p35* | 8 | 60-68 | 106-114 | 13 |
| 6C | *yw hsflp / w; UAS-dLMO / UAS-stg-RNAi; act>CD2>Gal4 UAS-GFP / +* | 8 | 60-68 | 106-114 | 13 |
| 6D | *yw hsflp / yw hsflp; UAS-dLMO / UAS-stg-RNAi; act>CD2>Gal4 UAS-GFP / UAS-p35* | 8 | 60-68 | 106-114 | 13 |
| 6F | Same as in 6B |  |  |  |  |
| 6G | Same as in 6D |  |  |  |  |
| S1A | *yw hsflp/(y)w; FRT<sup>f00878</sup> ubi-GFP / FRT<sup>f00878</sup>* | 4 | 42-46 | 66-70 | 30 |
| S1B | *yw hsflp/(y)w; FRT<sup>f00878</sup> ubi-GFP / FRT<sup>f00878</sup>* | 4 | 42-46 | 76-80 | 30 |
| S1C | *yw hsflp/(y)w; FRT<sup>f00878</sup> ubi-GFP / FRT<sup>f00878</sup>* | 4 | 42-46 | 86-90 | 30 |
| S2A | *yw hsflp / (y)w; FRT<sup>f00878</sup> / FRT<sup>f00878</sup> tub-Gal80; tub-Gal4 UAS-GFP / +* | 20 | 46-66 | - | 30 |
| S2B-G | *yw hsflp / (y)w; FRT<sup>f00878</sup> ap<sup>DG8</sup> / FRT<sup>f00878</sup> tub-Gal80; tub-Gal4 UAS-GFP / +* | 20 | 46-66 | - | 30 |
| S3A | *(w) hsflp / (y)w (hsflp); FRT<sup>f00878</sup> ap<sup>DG8</sup> ubi-GFP / FRT<sup>f00878</sup> tub-Gal80; tub-Gal4 UAS-mCherry/ UAS-p35* | 4 | 42-46 | 76-80 | 30 |
| S3B | *(w) hsflp / (y)w (hsflp); FRT<sup>f00878</sup> ap<sup>DG8</sup> ubi-GFP / FRT<sup>f00878</sup> tub-Gal80; tub-Gal4 UAS-mCherry/ UAS-p35* | 4 | 42-46 | 86-90 | 30 |
| S3C | *(w) hsflp / (y)w (hsflp); FRT<sup>f00878</sup> ap<sup>DG8</sup> ubi-GFP / FRT<sup>f00878</sup> tub-Gal80; tub-Gal4 UAS-mCherry/ UAS-p35* | 4 | 42-46 | 96-100 | 30 |
| S4A | *yw hsflp / w; IF or CyO / +; act>CD2>Gal4 UAS-GFP / MKRS* | 8 | 60-68 | 106-114 | 13 |
| S4B | *yw hsflp / hsflp; IF or CyO / +; act>CD2>Gal4 UAS-GFP / UAS-p35* | 8 | 60-68 | 106-114 | 13 |
| S4C | *yw hsflp / w; UAS-stgRNAi / +; act>CD2>Gal4 UAS-GFP / +* | 8 | 60-68 | 106-114 | 13 |
| S4D | *yw hsflp / yw hsflp; UAS-stgRNAi / +; act>CD2>Gal4 UAS-GFP / UAS-p35* | 8 | 60-68 | 106-114 | 13 |
| S4E | Same as in 6A |  |  |  |  |
| S4F | Same as in 6B |  |  |  |  |
| S4G | Same as in 6C |  |  |  |  |
| S4H | Same as in 6D |  |  |  |  |
| S5 | *yw hsflp /yw; FRT<sup>f00878</sup> ap<sup>DG8</sup> / FRT<sup>f00878</sup> tub-Gal80; tub-Gal4 UAS-GFP / +* | 6 | 61-67 | 108-114 | 30 |

